# Effect of resistant compartment on pathogen strategy in partially migratory populations

**DOI:** 10.1101/2024.06.14.599075

**Authors:** Cynthia Shao, Martha Torstenson, Allison K. Shaw

## Abstract

Migration, the recurring movement of animals between habitats, can exert pressures on the pathogens they host. Properties of host populations can determine pathogen strategy (e.g. virulence) to increase pathogen fitness. To study the effect of adding a resistant compartment on virulence evolution, we developed an SIRS model and examined the winning pathogen strategy across different rates of recovery and of immunity loss. We find that when hosts spend a relatively long time in the resistant class, a more virulent pathogen evolves. These results have implications in conservation of migratory animal populations afflicted by disease.

## Introduction

Understanding the factors that shape host-pathogen interactions is critical for both science broadly as well as for specific applications to animal, plant and human health challenges [1–3]. Two key aspects of host-pathogen interactions are virulence (here we use this specifically for disease-induced host mortality; [4]) and transmission rate.

Furthermore, there is often a tradeoff between transmission and virulence: high transmission rates come at the cost of increased host mortality, reducing the lifespan of the host and decreasing opportunity to spread, while pathogens that reproduce more slowly can do so at lower cost to the host, which increases the timespan in which a pathogen can spread but results in a lower transmission rate [5]. More specifically, the relationship between virulence and transmission is often convex where the curve of transmission as a function of mortality is concave down [6, 7].

One factor that can shape pathogen transmission and virulence is host movement.

Both theoretical and empirical work suggests that virulence should increase when infection can occur across long distances (i.e., as host movement increases) while parasites should be less virulent when transmission is local [8, 9]. However, theory also predicts that virulence should increase with natural host mortality, since this tends to decrease the duration of infection [10]. Since movement is often costly [11], increased movement could also lead to decreased virulence. A similar tension has been found in studying pathogen virulence in spreading host populations. When the only effect of parasite infection was to decrease host survival, the most virulent parasites were favored [12, 13]. However, if parasite infection simultaneously reduced host movement, lower virulence evolved [13]. However, the bulk of our knowledge of host movement comes from host dispersal, which is only one form of movement. In contrast, relatively less is known about how other forms of host movements like seasonal migration (recurring movement of animals between habitats) shape transmission and virulence.

Although much of the work exploring host-pathogen interactions in migratory species has focused on how pathogens shape host migration [14], we are increasingly learning about how host seasonal migration can shape pathogen virulence [15]. Recent theory shows that sedendary and migratory hosts can favor different pathogen virulence strategies [16]. Furthermore, properties of host populations can determine which pathogen strategies lead to the highest pathogen fitness [17]. For example, both the degree of tolerance that hosts have to pathogens as well as host pace of life can shape pathogen strategy [16].

However, this theory is based on only one of the possible types of host-pathogen systems [18, 19], with other yet to be explored. Specifically, the above model was an SIS (susceptible - infected - susceptible) compartmental model, based on a host-pathogen system where hosts that recover from infection can be immediately reinfected (i.e., no long-lasting immunity). In contrast, many pathogens are better described by an SIR (susceptible – infected - resistant) model where recovered hosts gain immunity in the resistant compartment [20]. Finally, in an SIRS (susceptible - infected - resistant - susceptible) model, resistant individuals do not have long lasting immunity and can move back into the susceptible compartment to possible be infected again. Since recovered and resistant individuals cannot immediately be re-infected, we expect that including a resistant compartment can affect pathogen evolution and virulence.

Whether or not immunity is acquired is an especially important consideration for migratory species. Migrants often come into contact with novel parasites during their journey [21] which can lead to migrants accumulating a greater diversity of parasites [22]. At the same time, because migration can be quite energetically demanding, migrants may be favored to reduce immune function during migration [23].

Here, we develop a model to understand how the presence of a resistant compartment affects pathogen evolution in a host population where some individuals migrate and others do not (partial migration; [24]). Specifically, we developed an SIRS model that included pathogen strategies of different virulence levels and transmission rates. We quantified how the best pathogen strategy varies with the rate of immunity loss and other key model parameters.

## 1 Methods and Model

### 1.1 Overview

The model investigates infection dynamics and pathogen strategy with an SIRS model. The host population consists of both migrants and residents to study how differences between the two groups (e.g. cost of migration, differing recovery rates during part of the year) affect infection dynamics. We investigate pathogen strategy by creating three strains of the infection with different virulence levels and simulated the number of individuals in each compartment of the SIRS model for 2000 years. At the end of each simulation, we determined which pathogen strategy was most common (i.e., most successful). We ran simulations across different values of the infected cost of migration, immunity loss rate, and recovery rate in habitat 1, in order to better understand the relationship between pathogen strategy and host migration strategy.

### 1.2 Compartments

We created an SIRS model with three infection classes (Fig 1). Infection class 1 was the least virulent, infection class 2 was intermediately virulent, and infection class 3 was the most virulent. These three infection classes provide a sufficient range of virulence levels to understand qualitatively how increasing virulence affects the behavior of the model. The host population was split into migrants and residents. There are 10 total classes in the model: susceptible residents (*S*_*r*_), class 1 infected residents 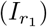, class 2 infected residents 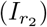, class 3 infected residents 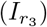, resistant residents (*R*_*r*_), susceptible migrants (*S*_*m*_), class 1 infected migrants 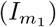, class 2 infected residents 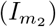, class 3 infected residents 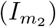, and resistant migrants (*R*_*m*_). Each compartment started with an initial population of 100 individuals. Migrants can move between the migrant compartments but cannot become residents and vice versa. However, migrants and residents affect each other through both pathogen transmission and density dependence (which is based on the total population of migrants and residents).

**Fig 1.**
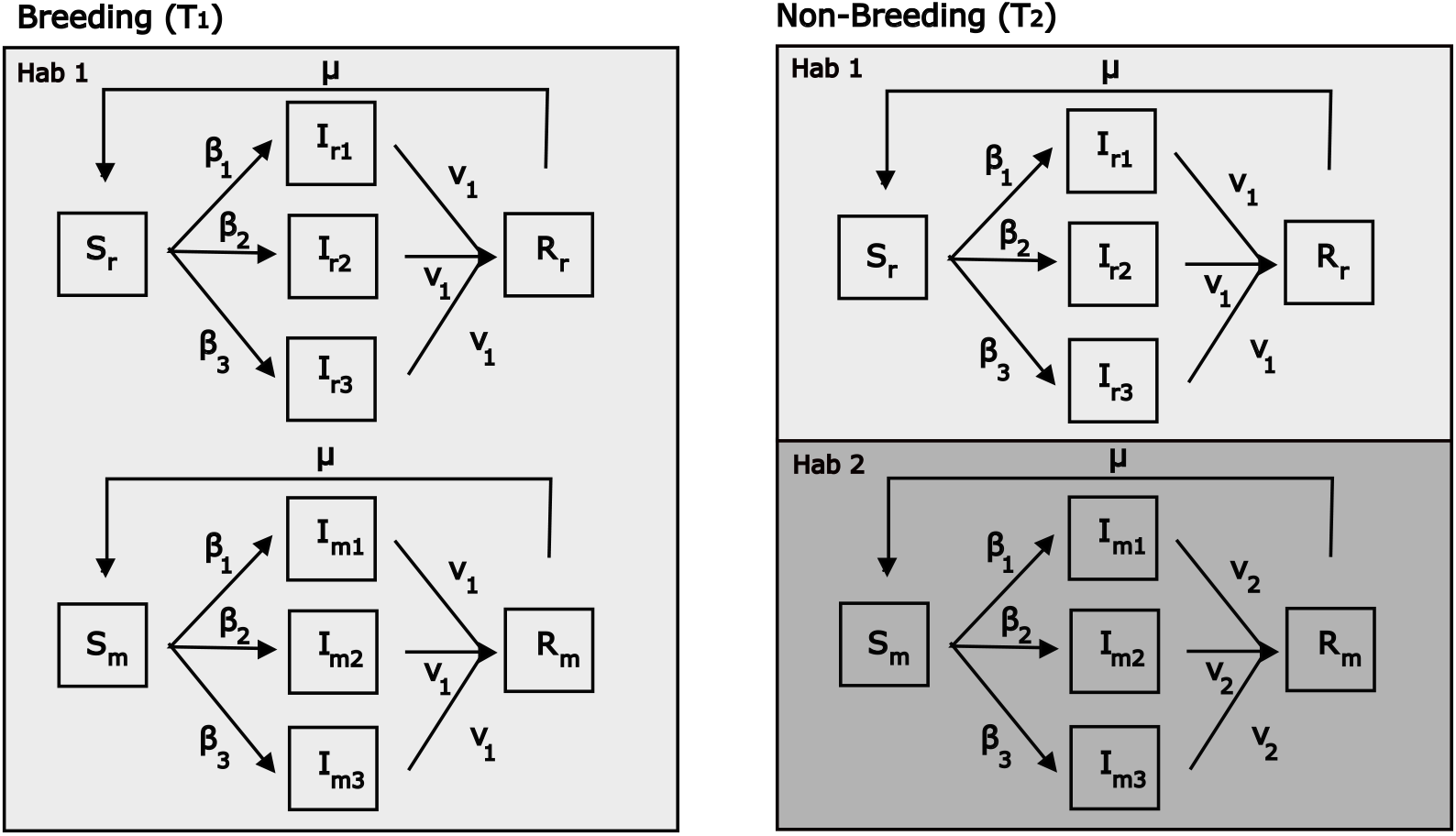
Model schematic. Migrants move between Habitat #1 and Habitat #2 while residents stay in Habitat #1 year long. Recovery rate is *ν*_1_ in Habitat #1 and *ν*_2_ in Habitat #2. See Table 1 for parameter values.

### 1.3 Transmission Rate and Virulence Trade-off

Pathogens often experience a trade-off between transmission rate and virulence, as the cost of increasing replication within the host (associated with more transmission) is increased host mortality. Transmission rate and virulence have a convex relationship [6]. Thus, as a pathogen’s virulence increases, the marginal increase in transmission associated with a given increase in virulence lowers. Pathogens with low transmission rates thus have lower virulence than pathogens with high transmission rates. Thus, low transmission rate pathogens keep their hosts alive for longer and have a longer amount of time to transmit to new hosts. A high virulence pathogen would infect individuals at a higher rate than the low virulence pathogen but would kill its host more quickly and thus has less time to proliferate and infect another host. In our model we describe transmission rate as

**Table 1.**
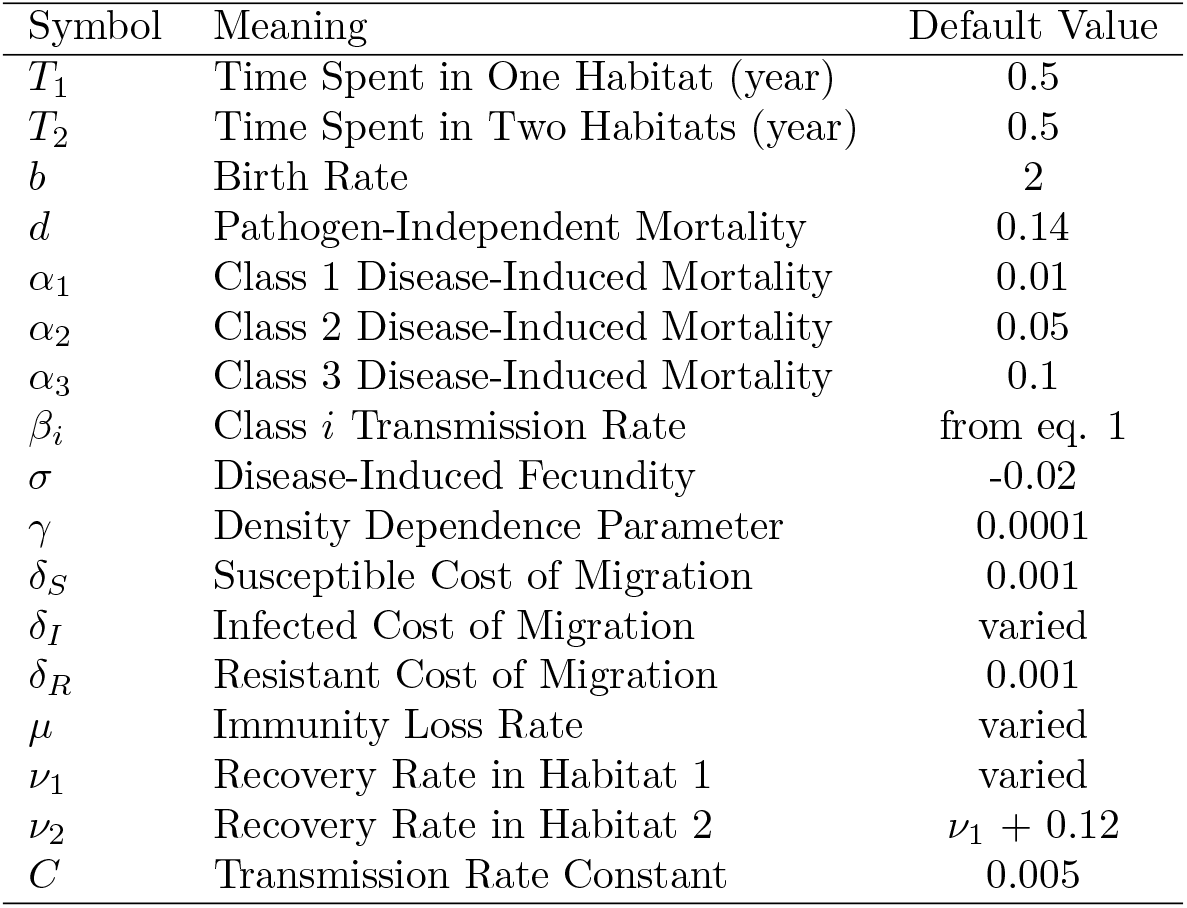
Model symbols, meanings, and default values.

| Symbol | Meaning | Default Value |
| --- | --- | --- |
| $T_1$ | Time Spent in One Habitat (year) | 0.5 |
| $T_2$ | Time Spent in Two Habitats (year) | 0.5 |
| $b$ | Birth Rate | 2 |
| $d$ | Pathogen-Independent Mortality | 0.14 |
| $\alpha_1$ | Class 1 Disease-Induced Mortality | 0.01 |
| $\alpha_2$ | Class 2 Disease-Induced Mortality | 0.05 |
| $\alpha_3$ | Class 3 Disease-Induced Mortality | 0.1 |
| $\beta_i$ | Class $i$ Transmission Rate | from eq. 1 |
| $\sigma$ | Disease-Induced Fecundity | -0.02 |
| $\gamma$ | Density Dependence Parameter | 0.0001 |
| $\delta_S$ | Susceptible Cost of Migration | 0.001 |
| $\delta_I$ | Infected Cost of Migration | varied |
| $\delta_R$ | Resistant Cost of Migration | 0.001 |
| $\mu$ | Immunity Loss Rate | varied |
| $\nu_1$ | Recovery Rate in Habitat 1 | varied |
| $\nu_2$ | Recovery Rate in Habitat 2 | $\nu_1 + 0.12$ |
| $C$ | Transmission Rate Constant | 0.005 |

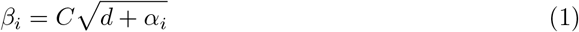

where *d* is the pathogen-independent mortality rate (the rate at which individuals die naturally) and *α*_*i*_ is the disease-induced mortality rate of pathogen class *i* (the added mortality from being infected), and *C* is a constant. We set *d* to 0.14, so that there was some mortality but the population was able to grow and persist in the absence of the pathogen. We set *C* to 0.005 to ensure a reasonable transmission rate where susceptible individuals were infected at a rate that would not overwhelm the population with infected individuals. Each infected class had a unique transmission rate dependent on the virulence level of the class. Transmission rate was the same for migrants and residents infected by the same infection class.

### 1.4 Loss of Immunity

We set up the model so that it could be run as either an SIR model (setting parameter *µ* = 0 defined below) or as an SIRS model (setting parameter 0 *< µ <* 1). For *µ* = 0, the model is effectively an SIR model with no movement from the resistant compartment to the susceptible compartment except for births. For *µ >* 0, some percentage of the resistant compartment loses resistance and moves back into the susceptible compartment in an SIRS model. As *µ* increases, individuals move out of the resistant compartment more quickly, and in the extreme the model effectively becomes an SIS model.

### 1.5 Evolutionarily Stable Strategy

In conflict situations, an evolutionarily stable strategy (ESS) is a strategy adopted by the majority of the population such that another competing strategy is unable to confer higher reproductive fitness [25]. In our model, pathogen strategies with different virulence levels are competing to be the ‘winning’ strategy within the host population population. We explored under which conditions each pathogen strategy wins. When an initial host population includes equal proportions of individuals infected with each pathogen class, the ‘winning’ pathogen strategy corresponds to the infection compartment that makes up the largest proportion of the final population. The winning strategy is considered the ESS. This often means that the other two pathogen strategies have population sizes that are ∼0.

### 1.6 Partial Migration

We set up the host population with migrant and resident individuals (i.e, with the potential for partial migration). During part of the year (*T*_1_), migrants and residents are in the same habitat and can infect each other. This is also the period of the year where births occurred. During the remainder of the year (*T*_2_), residents stay in the same habitat while migrants move to a separate habitat and return at the end of *T*_2_. We assume *T*_1_ + *T*_2_ = 1, i.e., migration is instantaneous in that we do not explicitly model infection dynamics during transit. When they are in separate habitats, residents can only be infected by residents and migrants can only be infected by migrants.

Transmission rates and disease-induced mortality rates were the same between residents and migrants infected with the same class of pathogen. Recovery rates (*ν*_2_), the movement from an infection compartment to a resistant compartment, were higher for migrants than residents during *T*_2_ to simulate migratory recovery. Recovery rate is *ν*_1_ for individuals in Habitat #1 and *ν*_2_ in Habitat #2. At the end of each simulation, we determined whether there were more migrants or residents.

### 1.7 Equations

We describe our model with two sets of equation, one for each time period *T*_1_ and *T*_2_. All parameter meanings and values are given in Table 1.

#### 1.7.1 One Habitat (*T*_1_)

During time *T*_1_ (when migrants and residents are both in the same habitat), the dynamics of susceptible residents is given by

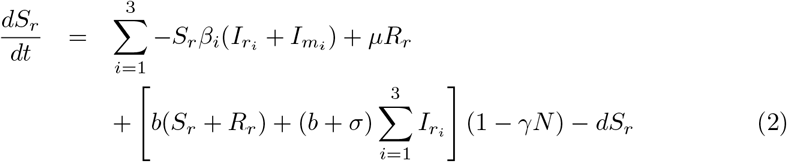

and the dynamics of susceptible migrants by

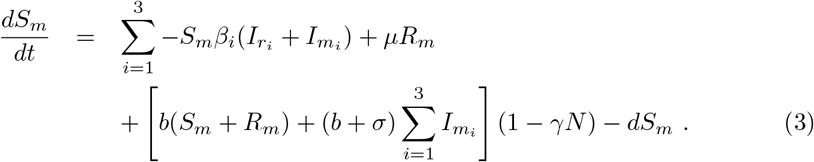

The first term in each equation describes how individuals move out of the susceptible compartment (to the *I* compartments) by becoming infected according to transmission rate *β*_*i*_, given by Eqn. (1). Susceptible individuals can be infected by any infected individual (regardless of whether they are a migrant or resident). The second term describes how individuals move in to the susceptible compartment (from the *R* compartments) when immunity is lost, at rate *µ*. The third term describes reproduction where *b* is the density-independent birth rate, *σ* is the disease-induced fecundity, *γ* is the strength of density-dependence in reproduction, and *N* is the total host population size (summed across both migrants and residents). We kept the birth rate (*b*) constant at 2, which was high enough to ensure population growth (compared to *d*) but not high enough to overwhelm the population with susceptible individuals. We set *γ* to 0.0001 to have a carrying capacity of 10,000 (to be relatively higher than our starting population of 1,000 individuals). As the total population *N* increases towards the carrying capacity, (1 − *γN*) decreases towards 0 and decreases the number of individuals born. Infected individuals produced less offspring due to the effects of the disease. All individuals born during *T*_1_ are susceptible. Finally, the last term in each equation describes pathogen-independent mortality at rate *d*.

The dynamics of infected residents and migrants are given by

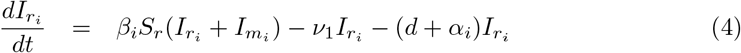

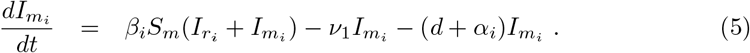

The first terms in each equation describe newly infected individuals. The second terms describes infected individuals that recover at rate *ν*_1_. The third term describes mortality, both from natural causes and from disease, at rate *d* + *α*_*i*_. We chose values for the disease-induced mortality rates (*α*_1_, *α*_2_, *α*_3_) that captured a wide range of possible detrimental effects of the pathogen while also allowing the population to reach steady state in a long enough time frame to investigate the model’s behavior.

The dynamics of resistant residents and migrants are given by

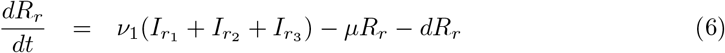

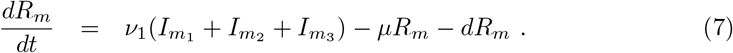

Here, the first terms describe individuals that have recovered from infection. The second term describes resistant individuals that lose immunity and move back to the susceptible compartments. The final term describes pathogen-independent mortality, as above.

#### 1.7.2 Two Habitats (*T*_2_)

During time *T*_2_ (when migrants and residents are in two separate habitats), the dynamics of susceptible residents and migrants is given by

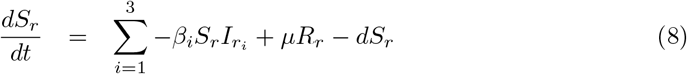

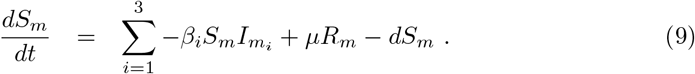

The dynamics are similar to those during *T*_1_ (Eqns. 2-3) except that residents are only infected by other residents, and migrants are only infected by other migrants, and no reproduction occurs. The dynamics of infected residents and migrants is given by

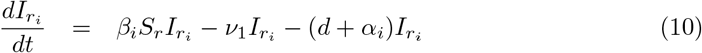

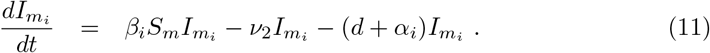

We assume that recovery rate is habitat-dependent, so infected residents recover at rate *ν*_1_ while infected migrants recovers at a higher rate *ν*_2_. We varied *ν*_1_) and set *ν*_2_ = *ν*_1_ + 0.12 to ensure a noticeable benefit for migrants who survive the process of migration. Finally, the dynamics of resistant residents and migrants are given by

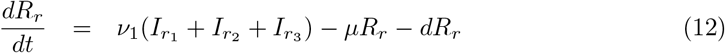

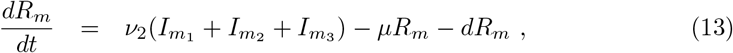

which is identical to the dynamics during *T*_1_ (Eqns. 6-7).

We assume that migrants experience a mortality cost each time they migrate. To capture this cost, twice a year (at the end of *T*_1_ and *T*_2_), we multiply the number of susceptible migrants by 1 − *δ*_*S*_, the number of infected migrants by 1 − *δ*_*I*_, and the number of resistant migrants by 1 − *δ*_*R*_, so the actual number of individuals is

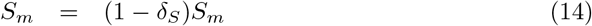

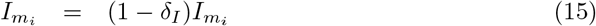

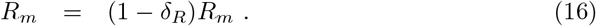

The susceptible and resistant cost of migration (*δ*_*S*_ and *δ*_*R*_) were set to 0.001 to ensure there was some detrimental consequence to migrating, but not enough to overly favor the residents.

### 1.8 Simulations

We started each simulation with 100 individuals in each of the 10 classes and ran it for 2000 years. We ran simulations under different values of the immunity loss rate (*µ*), the recovery rate in habitat 1 (*ν*_1_) and the infected cost of migration *δ*_*I*_ in order to explore interactions between migration dynamics and pathogen strategy. We kept the remaining parameters constant. For each simulation, we quantified the number of individuals by pathogen strategy and by host migration strategy.

## 2 Results

We can examine the output of each simulation run of our model in a number of ways. We tracked the total number of individuals of each of the 10 host types over time – the number of susceptible, infected (with each of the three pathogen types) and recovered individuals for each migrant and resident hosts (Fig 2a). We determined which pathogen strategy won by combining the number of migrant and resident hosts infected with each pathogen type; typically we found that a single pathogen strategy was much more abundant than the other two (Fig 2b). We also determined what happened to the host types by combining the number of hosts within each the migrant and host categories; again we typically found that a single host strategy (migrant or resident) was more abundant than the other (Fig 2c).

**Fig 2.**
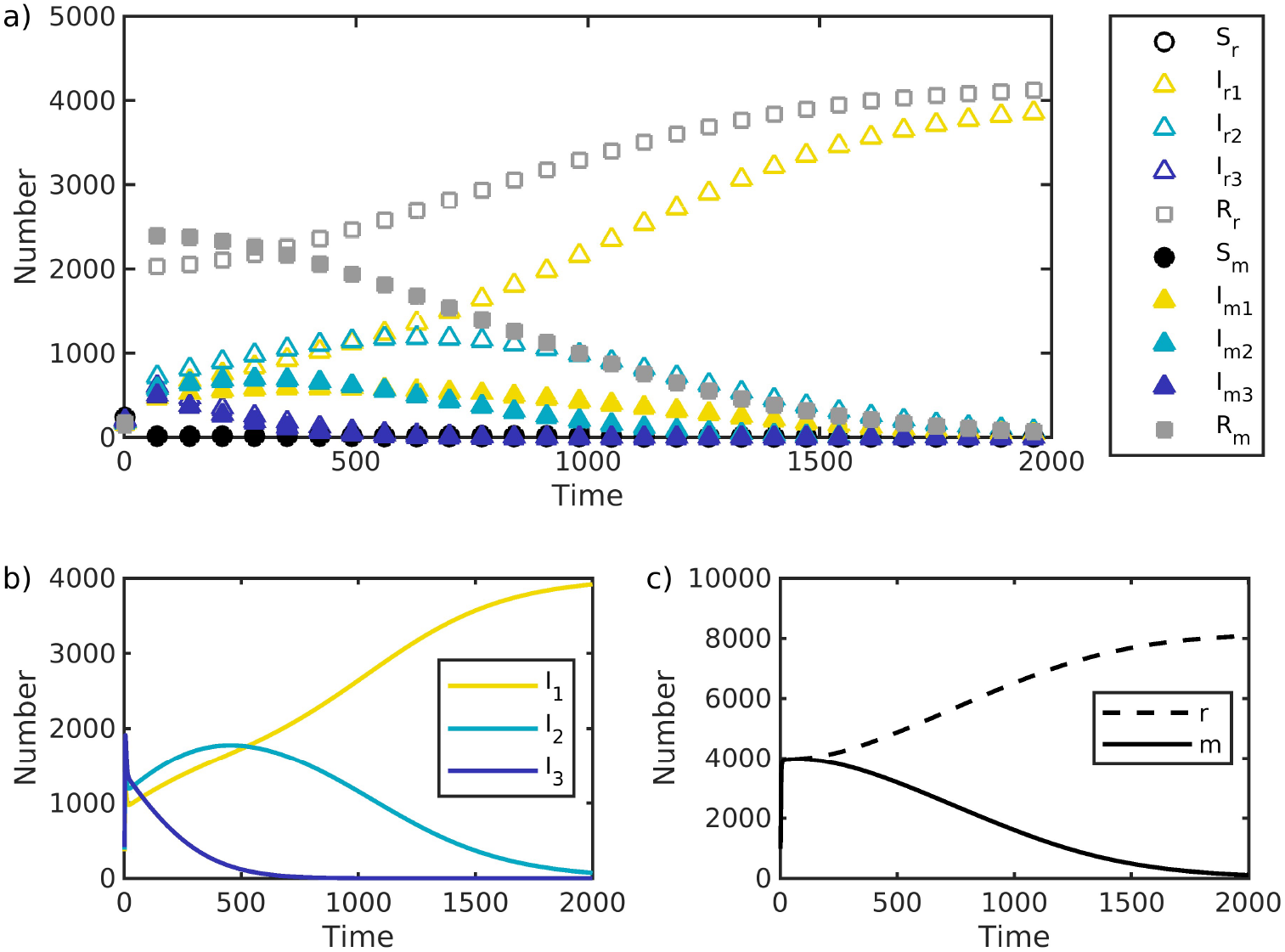
Example of a single simulation output showing the number of host individuals of each type over time for (a) all 10 types: *S* suceptible, *I* infected, and *R* recovered individuals: for *r* residents and *m* migrants, (b) the number of individuals by pathogen class (1, 2, or 3), (c) the number of individuals by host strategy (residents or migrants). Parameter values: *δ*_*S*_ = *δ*_*R*_ = 0.001, *δ*_*I*_ = 0.004, *ν*_1_ = 0.14, *ν*_2_ = *ν*_1_ + 0.12; and *µ* = 0. Other parameter values given in Table 1.

Our results show that there are cases where each of the three pathogen strategies can win, i.e. infect the most hosts in the population (Fig 3). Which pathogen strain did best in our model depended in large part on how quickly hosts recovered from infection. In particular, pathogen strategy 1 (which has the lowest virulence) was favored when recovery rate (*ν*_1_) was relatively low, while pathogen strategy 1 (which has the highest virulence) was favored when *ν*_1_ was high (Fig 3). Thus, increasing recovery favored an increase in pathogen virulence. To a lesser extent, the pathogen strategy that did best also depended on how quickly host immunity was lost, where faster rates of immunity loss (*µ*) favored slightly less virulent pathogens (Fig 3). These results generally hold across all values of *δ*_*I*_ (the cost of migration for infected individuals) that we considered (Fig 4).

**Fig 3.**
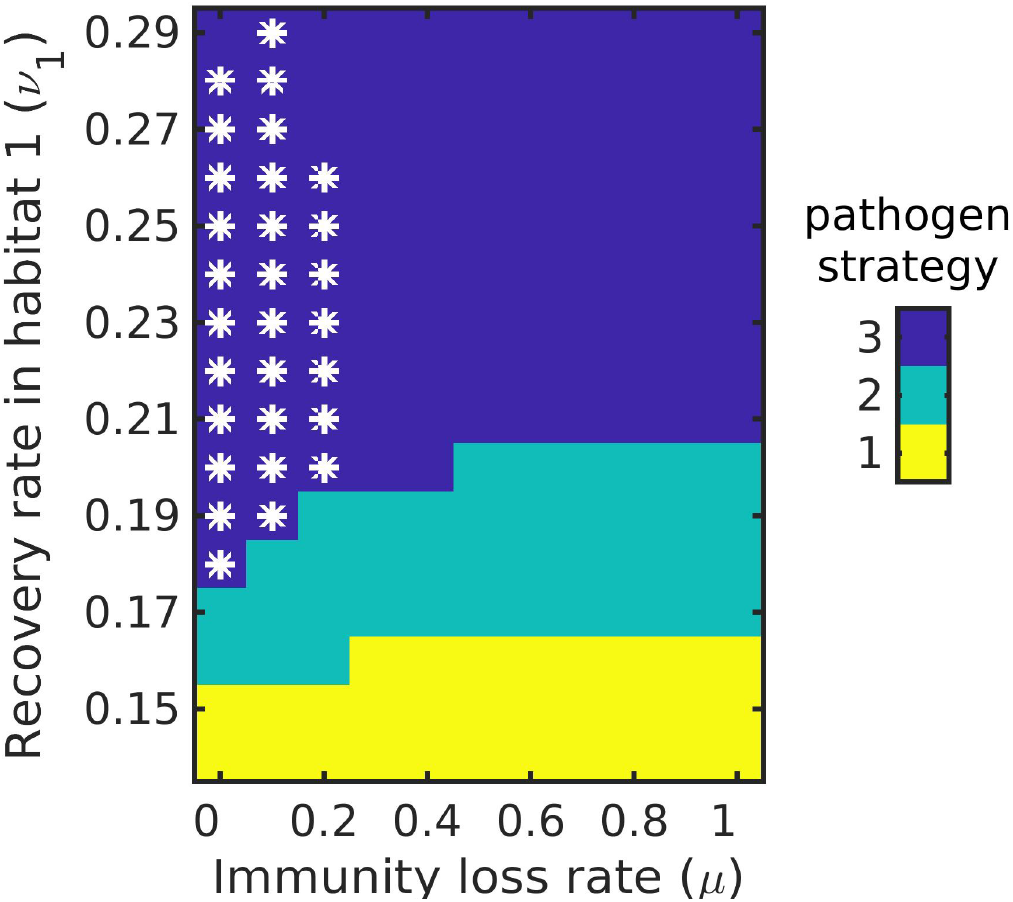
The pathogen class (1, 2 or 3) that wins as a function of the immunity loss rate (*µ*; x-axis) and recovery rate (*ν*_1_; y-axis). Parameter values: *δ*_*S*_ = *δ*_*R*_ = 0.001, *δ*_*I*_ = 0.004, *ν*_2_ = *ν*_1_ + 0.12; *µ* and *ν*_1_ were varied. Other parameter values given in Table 1. Asterisks indicate where there were more migrants than residents.

**Fig 4.**
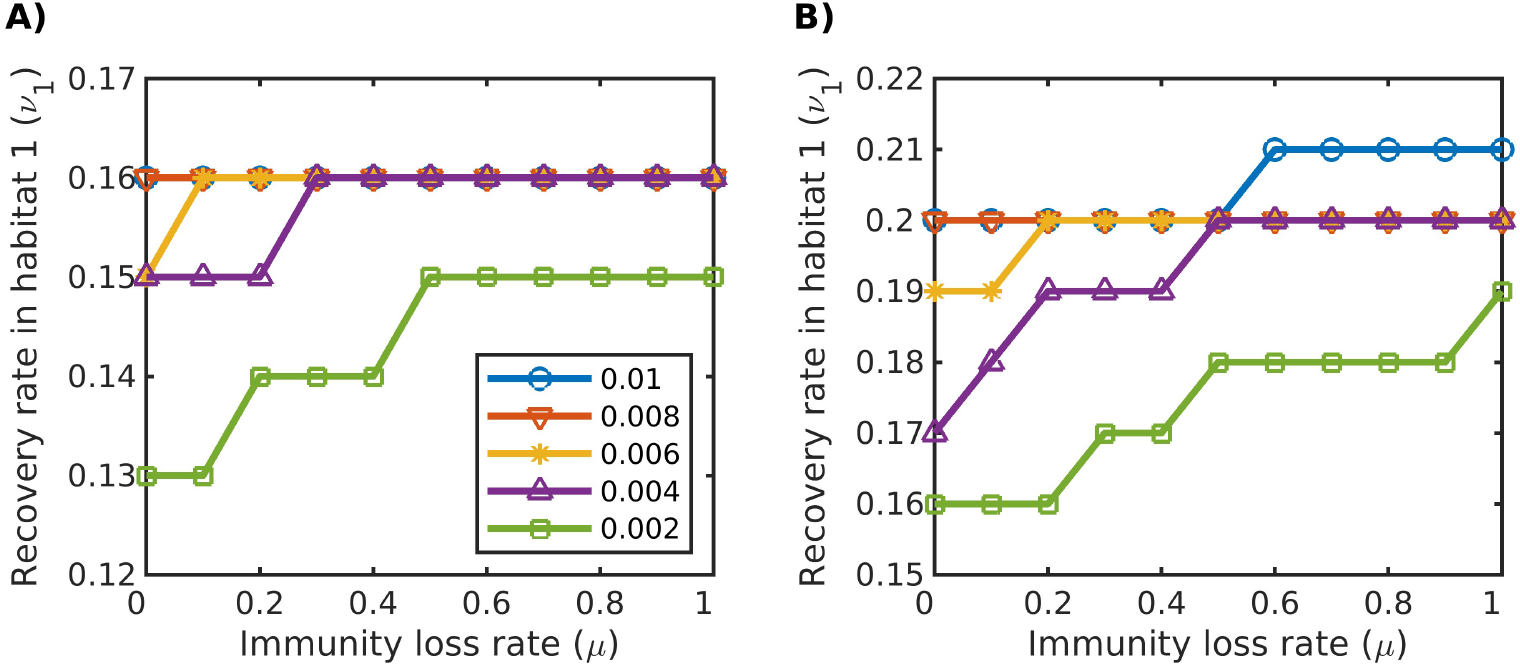
The recovery rate (*ν*_1_; y-axis) at which the winning infection class changes **A)** from class 1 to class 2, and **B)** from class 2 to class 3, as a function of the immunity loss rate (*µ*; x-axis), for different values of *δ*_*I*_ (colors), the infection cost of migration. Parameter values: *δ*_*S*_ = *δ*_*R*_ = 0.001, *ν*_2_ = *ν*_1_ + 0.12; *µ, ν*_1_, and *δ*_*I*_ were varied. Other parameter values given in Table 1.

### 2.1 Resistant compartment mediates interaction between recovery and immunity loss rates

To understand how the existence of a resistance class affects virulence, we can compare our results across different parameter values. For *ν*_1_ (no recovery) the model is effectively an SI model: individuals can get infected but never recover. This favors low pathogen virulence. For *ν*_1_ *>* 0 but *µ* = 0, the model is effectively an SIR model: individuals recover but never lose immunity. Here, the virulence of the pathogen strategy increases as *ν*_1_ increases. Finally, for *ν*_1_ *>* 0 and *µ >* 0, we have an SIRS model. By varying the relative values of *ν*_1_ and *µ*, we vary the average amount of time that hosts spend immune once they recover from infection. A high *µ* value and low *ν*_1_ value mean that hosts recovery slowly and lose immunity quickly, thus spending little time in the R compartment. In contrast a low *µ* value and high *ν*_1_ value mean that hosts recovery quickly and lose immunity slowly, thus spending a longer time in the R compartment. Looking diagonally across Fig 3, we can see that spending longer in the R compartment (moving from the lower right to the upper left) favors for increased pathogen virulence.

### 2.2 Interaction between migration and virulence

In our model, infected individuals face a trade-off when it comes to migration: infected individuals have a higher cost of migration than susceptible and resistant individuals, but gain the benefit of increased recovery while in the migratory habitat (since *ν*_2_ *> ν*_1_). Which pathogen strain did best in our model thus also depended in part on how costly migration was for infected individuals. We can plot the values of *ν*_1_ and *µ* where we see a switch from pathogen strategy 1 to pathogen strategy 2 being favored (Fig 4). Doing this, we see that when the cost of migration for infected individuals (*δ*_*I*_) is high, the switch from strategy 1 to strategy 2 occurs for higher values of *ν*_1_ and *µ* than if *δ*_*I*_ is low (Fig 4). In other words, as the infection-based cost of migration increases, less virulent pathogens are favored. Finally, we also typically found that there were more residents than migrants, except when *µ* was quite low (Fig 3). This indicates the point at which the benefit of increased recovery rate in the migratory habitat overcomes the cost of migration.

## Discussion

### Effect of host migration

Here, we developed a model to understand how the presence of a resistant compartment, combined with host migratory behavior, affected pathogen evolution. Our first finding was that host migration can indeed shape pathogen virulence. Specifically we found that a lower cost of migration (e.g. a lower *δ*_*I*_ in our model) increases pathogen virulence (Fig 4), presumably by increasing the proportion of migrants within the population.

This result parallels theory on other forms of host movement (i.e. dispersal). Past work finds that increased host dispersal favors pathogens that enter and exit the host quickly [26]. Similarly, past theory has found that populations with restricted spatial movement evolve lower virulence, and populations with greater connectivity evolve higher virulence [8, 27]. This result is also found empirically in bacteriophage systems, which found that less virulent strains are favored when movement is restricted [27].

The relative number of migratory and resident hosts also varied in our model. When the cost of migration for infected individuals is sufficiently low and the recovery rate for migrants is sufficiently high, migrants are able to win over the residents. This is likely because migrants are only able to maintain their population under circumstances with low cost of migration and high recovery. We found that conditions where the migrant population was greater than the resident population, were also conditions where a more virulent pathogen strain won (Fig 3). In contrast to this finding, studies on butterfly populations found that parasites from resident winter-breeding sources had the same virulence as parasites from migratory sources, and various other studies found that increased transmission opportunities featured increased virulence [28]. Theoretically, an increased proportion of migrants could lead to reduced virulence due to migratory escape and migratory culling [28]. In a sense, we see this result in our model as well – although we do not have migratory escape, we do have migratory culling (the degree to which *δ*_*I*_ is bigger than *δ*_*S*_ or *δ*_*R*_), as migratory culling increases (*δ*_*I*_ gets bigger), we see a shift towards less virulent pathogens (Fig 4), although this may be because hosts shift to becoming non-migratory.

### Effect of resistant compartment

Our second finding was that the presence of a resistant compartment–and how long hosts spent in that compartment–shaped pathogen virulence. In an SIR model (*µ* = 0), a pathogen with a high recovery rate may favor a high virulence strategy, as the pathogen will have less time to reproduce in the host and will aggressively use the host’s resources. A low recovery rate may favor a low virulence strategy, as the pathogen can stay in the host longer to proliferate and infect other possible hosts. However, in SIRS models where resistant individuals can lose immunity and move back into the susceptible compartment, the strength of this trend may be reduced, thought the direction of the effect remains consistent. If there is a steady supply of susceptible individuals to infect, the pathogen may favor a high virulence strategy regardless of recovery rate, as killing the host rapidly is less of a concern if there are many susceptible individuals to infect. We see this reflected in Fig 3 and Fig 4, where higher recovery rates favor higher virulence strains.

The rate of immunity loss, *µ*, moves individuals from the resistant compartment to the susceptible compartment. A higher *µ* means that a population has fewer resistant individuals and more susceptible individuals compared to a lower *µ*, where the population would consist of more resistant individuals and fewer susceptible individuals. A population with a smaller proportion of susceptible individuals could favor a less virulent strain, as slowly expending host resources to spread the pathogen would be preferable to killing the host quickly when there are not many individuals to infect at a given point in time. Waiting in the host means the pathogen can infect more individuals. By increasing the number of susceptible individuals when having resistant individuals lose immunity, the size of the susceptible compartment increases. However, pathogen strategy itself also impacts the number of susceptible individuals present.

Highly virulent pathogens reduce the number of resistant hosts (i.e. by killing off infected hosts before they recover), which otherwise would block transmission [29]. When recovery and immunity loss are both high, the most virulent pathogen strain wins (Fig 3), as the pathogen has a continuous supply of susceptible hosts to infect and the infected individuals do not stay infected for long, meaning the pathogen will rapidly exploit the host’s resources.

### Parallels to past work

We can draw a number of parallels between our findings here and past work about the evolution of pathogen virulence. First, shorter infection durations should typically lead to more virulent pathogen strategies. In this paper, we found that shorter infection durations can be caused by higher recovery rates. In past work, we have shown that shorter duration caused by higher mortality also leads to a more virulent pathogen strategy [16]. Second, hosts that are better adapted to cope with pathogen infection can lead to higher virulence. This can occur e.g. via hosts maintaining immunity for longer (lower immunity lost *µ* in our model), or can occur if hosts tolerate infection more (i.e. experience lower mortality) [16]. Third, the abundance of susceptible individuals available to become infected has a stronger effect on virulence than loss of infected individuals. Here we see this result in that loss of immunity by recovered individuals matters more than recovery of infected individuals (Fig 4). In past work we found that fecundity matters more than mortality [16].

### Future Directions

Future work could explore different sources of density-dependence. A mortality-based density dependent model could provide further insight into population dynamics. For fecundity-based density dependence (as implemented in this model), the birth rate decreases when the population nears carrying capacity and mortality rate is held steady. For mortality-based density dependence, the mortality rate increases when the population nears carrying capacity and birth rate is held steady. For a fecundity-based model, if resistant individuals make up the majority of the population and the population nears the carrying capacity, then the number of susceptible individuals would decrease (due to a decreasing birth rate). On the other hand, a mortality-based model would increase mortality rate among all compartments and the rate at which new susceptible individuals enter the population through fecundity will not be affected by host population density. Thus, differing results between models that employ a fecundity-based density dependence approach or a mortality-based density dependence approach may be possible.

## Conclusion

Our results have implications for conservation in partially migratory animal populations to understand disease dynamics. Building understanding of how pathogen resistance and lost resistance can affect virulence evolution in a partially migratory species can aid in animal populations experiencing a disease outbreak.

## Author Contributions

AKS conceived the idea and secured funding; MT and AKS formulated the research goals; CS coded the model, CS developed and analysed the model in consultation with MT; CS wrote the original draft; and CS, MT and AKS edited the final draft.

## Code availability

Code will be made available through Zenodo upon manuscript acceptance.

## Acknowledgements

We thank the Shaw Lab for helpful discussions. This material is based in part upon work supported by an NSF Research Experience for Undergraduates (REU) component of NSF Grant No. DEB-1947406.

## References

1. Ashander J, Krkošek M, Lewis MA. Aquaculture-induced changes to dynamics of a migratory host and specialist parasite: a case study of pink salmon and sea lice. Theoretical Ecology. 2011;5(2):231–252. doi:10.1007/s12080-011-0122-4.

2. Van Hemert C, Pearce JM, Handel CM. Wildlife health in a rapidly changing North: focus on avian disease. Frontiers in Ecology and the Environment. 2014;12(10):548–556. doi:10.1890/130291.

3. Mitchell CE, Blumenthal D, Jarošík V, Puckett EE, Pyšek P. Controls on pathogen species richness in plants’ introduced and native ranges: roles of residence time, range size and host traits. Ecology Letters. 2010;13(12):1525–1535. doi:10.1111/j.1461-0248.2010.01543.x.

4. Anderson RM, May RM. Coevolution of hosts and parasites. Parasitology. 1982;85(02):411–426.

5. Acevedo MA, Dillemuth FP, Flick AJ, Faldyn MJ, Elderd BD. Virulence-driven trade-offs in disease transmission: A meta-analysis*. Evolution. 2019;73(4):636–647. doi:10.1111/evo.13692.

6. Alizon S, van Baalen M. Emergence of a convex trade-off between transmission and virulence. The American Naturalist. 2005;165(6):E155–E167. doi:10.1086/430053.

7. Lipsitch M, Moxon ER. Virulence and transmissibility of pathogens: what is the relationship? Trends in Microbiology. 1997;5(1):31–37. doi:10.1016/S0966-842X(97)81772-6.

8. Boots M, Sasaki A. ‘Small worlds’ and the evolution of virulence: infection occurs locally and at a distance. Proceedings of the Royal Society of London B: Biological Sciences. 1999;266(1432):1933–1938.

9. Kerr B, Neuhauser C, Bohannan BJM, Dean AM. Local migration promotes competitive restraint in a host–pathogen ‘tragedy of the commons’. Nature. 2006;442(7098):75–78. doi:10.1038/nature04864.

10. Restif O, Hochberg ME, Koella JC. Virulence and age at reproduction: new insights into host-parasite coevolution. Journal of Evolutionary Biology. 2001;14(6):967–979. doi:10.1046/j.1420-9101.2001.00355.x.

11. Bonte D, Van Dyck H, Bullock JM, Coulon A, Delgado MdM, Gibbs M, et al. Costs of dispersal. Biological Reviews. 2012;87:290–312. doi:10.1111/j.1469-185X.2011.00201.x.

12. Griette Q, Raoul G, Gandon S. Virulence evolution at the front line of spreading epidemics. Evolution. 2015;69(11):2810–2819. doi:10.1111/evo.12781.

13. Osnas EE, Hurtado PJ, Dobson AP. Evolution of pathogen virulence across space during an epidemic. The American Naturalist. 2015;185(3):332–342. doi:10.1086/679734.

14. Binning SA, Craft ME, Zuk M, Shaw AK. How to study parasites and host migration: a roadmap for empiricists. Biological Reviews. 2022;97(3):1161–1178. doi:10.1111/brv.12835.

15. Poulin R, de Angeli Dutra D. Animal migrations and parasitism: reciprocal effects within a unified framework. Biological Reviews. 2021;96(4):1331–1348.

16. Torstenson M, Shaw AK. Pathogen evolution following spillover from a resident to a migrant host population depends on interactions between host pace of life and tolerance to infection. Journal of Animal Ecology. 2024;93(4):475–487. doi:10.1111/1365-2656.14075.

17. Ewald PW. The evolution of virulence: A unifying link between parasitology and ecology. The Journal of Parasitology. 1995;81(5):659. doi:10.2307/3283951.

18. Ross R. An application of the theory of probabilities to the study of a priori pathometry.—Part I. Proceedings of the Royal Society of London Series A, Containing Papers of a Mathematical and Physical Character. 1916;92(638):204–230. doi:10.1098/rspa.1916.0007.

19. Batista AM, De Souza Slt, Iarosz KC, Almeida ACL, Szezech Jr JD, Gabrick EC, et al. Simulation of deterministic compartmental models for infectious diseases dynamics. Revista Brasileira de Ensino de Física. 2021;43:e20210171. doi:10.1590/1806-9126-rbef-2021-0171.

20. Kermack WO, McKendrick A. A contribution to the mathematical theory of epidemics. Proceedings of the Royal Society A. 1927;115:700–721.

21. Jenkins T, Thomas GH, Hellgren O, Owens IP. Migratory behavior of birds affects their coevolutionary relationship with blood parasites. Evolution. 2012;66(3):740–751. doi:10.5061/dryad.qr8v5f4v.

22. Koprivnikar J, Leung TLF. Flying with diverse passengers: greater richness of parasitic nematodes in migratory birds. Oikos. 2015;124(4):399–405. doi:10.1111/oik.01799.

23. Hegemann A, Fudickar AM, Nilsson J A physiological perspective on the ecology and evolution of partial migration. Journal of Ornithology. 2019;160(3):893–905. doi:10.1007/s10336-019-01648-9.

24. Chapman BB, Brönmark C, Nilsson J Hansson LA. The ecology and evolution of partial migration. Oikos. 2011;120:1764–1775.

25. Maynard Smith J, Price GR. The logic of animal conflict. Nature. 1973;246:15–18. doi:10.1038/246015a0.

26. Eshelman CM, Vouk R, Stewart JL, Halsne E, Lindsey HA, Schneider S, et al. Unrestricted migration favours virulent pathogens in experimental metapopulations: evolutionary genetics of a rapacious life history. Philosophical Transactions of the Royal Society B: Biological Sciences. 2010;365(1552):2503–2513. doi:10.1098/rstb.2010.0066.

27. Cressler CE, McLeod DV, Rozins C, van den Hoogen J, Day T. The adaptive evolution of virulence: a review of theoretical predictions and empirical tests. Parasitology. 2016;143(7):915–930. doi:10.1017/S003118201500092X.

28. Satterfield DA, Maerz JC, Altizer S. Loss of migratory behaviour increases infection risk for a butterfly host. Proceedings of the Royal Society B: Biological Sciences. 2015;282(1801):20141734–20141734. doi:10.1098/rspb.2014.1734.

29. Boots M, Hudson PJ, Sasaki A. Large shifts in pathogen virulence relate to host population structure. Science. 2004;303:842–844.

